# Genome Repository of Oiled Systems (GROS): an interactive and searchable database that expands the catalogued diversity of crude oil-associated microbes

**DOI:** 10.1101/838573

**Authors:** Smruthi Karthikeyan, Luis M. Rodriguez-R, Patrick Heritier-Robbins, Janet K. Hatt, Markus Huettel, Joel E. Kostka, Konstantinos T. Konstantinidis

**Author notes:** Address correspondence to Konstantinos Konstantinidis.

## Abstract

Indigenous microbial communities ultimately control the fate of petroleum hydrocarbons (PHCs) that enters the natural environment through natural seeps or accidental oil spills, but the interactions among microbes and with their chemical environment during oil biodegradation are highly complex and poorly understood. Genome-resolved metagenomics have the potential to help in unraveling these complex interactions. However, the lack of a comprehensive database that integrates existing genomic/metagenomic data from oiled environments with physicochemical parameters known to regulate the fate of PHCs currently limits data analysis and interpretations. Here, we present a curated, comprehensive, and searchable database that documents microbial populations in oiled ecosystems on a global scale, along with underlying physicochemical data, geocoded via GIS to reveal geographic distribution patterns of the populations. Analysis of the ~2,000 metagenome-assembled genomes (MAGs) available in the database revealed strong ecological niche specialization within habitats *e.g.*, specialization to coastal sediments vs. water-column vs. deep-sea sediments. Over 95% of the recovered MAGs represented novel and uncultured species underscoring the limited representation of cultured organisms from oil-contaminated and oil reservoir ecosystems. The majority of MAGs linked to oiled ecosystems are members of the rare biosphere in non-oiled samples, except for the Gulf of Mexico (GoM) which appears to be primed for oil biodegradation. GROS should facilitate future work toward a more predictive understanding of the microbial taxa and their activities that control the fate of oil spills as well as serve as a model approach for building similar resources for additional environmental processes and omic data of interest.

## MAIN

Oil spills have pronounced impacts on natural ecosystems and the microbial community dynamics following such spills have only recently been documented, as best exemplified by the Deepwater Horizon (DWH) discharge in the GoM, the largest accidental marine oil spill in history. Biodegradation mediated by a complex network of microorganisms is the ultimate fate of the majority of petroleum hydrocarbons (PHCs) that enter the natural environment from accidental discharges ^1–4^. The microbial interactions that ultimately dictate the fate of oil are nonetheless highly complex and remain poorly understood ^5^. Reconstruction of metagenome-assembled population genomes (MAGs) from metagenomic data ^6,7^ enables the recovery of the functional potential of microbial consortia that are associated with critical ecosystem processes such as carbon cycling ^8,9^. Consequently, this approach can help to anchor microbial genomes to their biogeochemical functions and ecological niches, and thus advance understanding of the complex interactions that dictate the ecosystem functions facilitated by microorganisms. Public repositories host a plethora of sequence data from a diverse array of oiled environments, especially raw-reads or assembled MAGs. However, lack of environmental context in the form of *in-situ* physicochemical data that can be easily searched has severely hampered the usefulness and interpretations of the omics data. For instance, a multitude of spatio-temporal “omics” data following the DWH oil spill in the GoM, the first major oil spill for which a lot of omics datasets became available and revealed a short-term decrease in microbial community diversity accompanied by an increase in the functional repertoire ^10^. Moreover, specific taxon succession patterns were observed across spatial and temporal scales, which in turn, paralleled the chemical evolution of PHCs ^10–12^. However, the universal applicability of these patterns and the impacts of oil on ecosystem functions as well as the identification of “bioindicator” taxa require further study for use in emergency response efforts.

Specifically, the dynamics of key members of the rare biosphere across the range of environmental parameters observed in oiled marine ecosystems or habitats (e.g., coastal sediments vs. water-column vs. deep-sea sediments) at a global scale requires confirmation with more robust datasets. Such information could validate, for example, universal biomarker taxa and/or genes as indicators for the different stages of oil biodegradation (e.g., taxa that are dominant at early, mid vs. late stages) and thus, help in monitoring the fate of the spills and subsequent ecosystem recovery. Currently, no resource exists to taxonomically classify and provide biogeographic distribution and *in-situ* physicochemical or environmental context for a given MAG to enable these lines of research. The availability of associated *in-situ* physicochemical data could also help to disentangle the complex interactions that govern microbial community structuring and functioning post disturbance as well as to aid in culture/isolate novel keystone taxa. For instance, our own recent efforts recovered the genome of a population that strongly responded to oil contamination in GoM beach sands that was subsequently shown to comprise 20-30% of the total microbial community in oiled coastal sediments worldwide, and guided the isolation of a representative strain, *Ca.* Macondimonas diazotrophica (Karthikeyan et al., 2019). *Ca*. M. diazotrophica was demonstrated to mediate nitrogen fixation along with hydrocarbon degradation, a combination of traits that likely provide an important ecological advantage in oiled environments that are often nutrient-limited ^13^.

In the present study, we curated the “**Genome Repository of Oiled Systems**” (“GROS”), a comprehensive MAG and single-amplified genome (SAG) reference database that is provided as an independent and searchable project through the Microbial Genomes Atlas webserver (MiGA) ^14^. Since the DWH discharge represents the first major oil spill occurring after the development of next generation sequencing technologies, GROS is currently dominated by DWH data as well as MAGs binned from publicly available data from other oil-impacted environments, such as natural oil seeps and lab incubation or enrichment studies designed to mimic oil spills under near *in situ* conditions (Suppl. Table). The interactive graphical interface allows for browsing the MAGs/SAGs/isolate genomes in the GROS project and their associated metadata. GROS also enables users to query their isolate or MAG genome sequence (partial or complete) against its reference MAGs and SAGs to identify if the queried genome represents a new taxon (species) or is another member of taxa that is already represented among the reference MAGs. For query genomes of the same species as a reference MAG, the GROS project can be used to assess the global biogeographical distribution of the reference MAG on an interactive map and obtain information about the habitats/samples where the latter has been recovered and its relative abundance within these samples (when a corresponding metagenome is available). Therefore, the resulting data can help determine the taxonomic uniqueness of a queried genome, its relative *in-situ* abundance, and the extent of association with oiled samples based on existing data. The taxonomic classification of reference or user-provided query genomes is performed by MiGA as described previously based on the ANI/AAI concept ^14^. Incorporation of embedded GIS-based “layers” as part of the metadata description of each reference MAG provides an interactive interface that enables filtering based on criteria including location, habitat, taxonomy, date, nutrient concentrations, and hydrocarbon data, among other parameters (Fig. 1). The incorporation of the MAGs and their associated metadata as GIS layers enables overlaying additional *in-situ* physicochemical data available through the Environmental Response Management Application, ERMA (https://response.restoration.noaa.gov/gulf-mexico-erma), an online mapping tool which contains real-time data collected from the GoM.

**Fig. 1:**
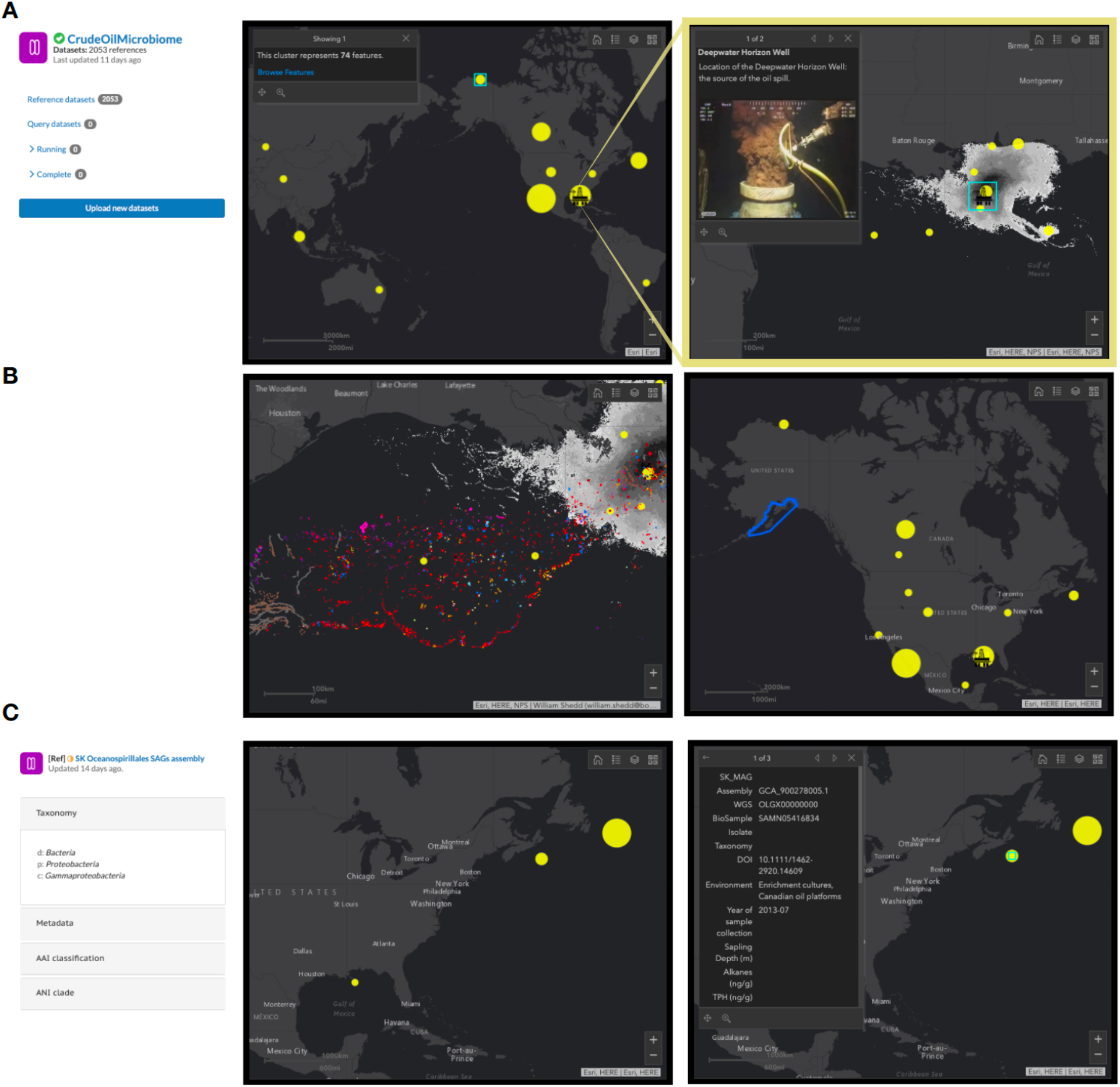
The graphical output from the GROS webserver. **A.** The map shows the distribution of all MAGs present in the curated COM database. The circles are proportional to the abundance of MAGs associated with each location. The top right panel shows only the MAGs recovered from the Gulf of Mexico (GoM) and the grey area denotes the cumulative days of oiling following the DWH accident along with the location of the Macondo MC252 wellhead. The interactive map allows enabling/disabling any or all of the other layers; namely, oil spill boundaries from previous oil spills, natural seep locations based on seismic surveys, as well as filtering the MAGs based on user-defined genomic criteria such as taxonomic name and genome completeness level. **B. Enabling additional layers.** The map layer showing the natural seep locations in the GoM region has been enabled. Bottom right: The modified map when the layer containing previous oil spill boundaries (in this instance, the Exxon-Valdez oil-spill) have been included. **C**. **Assessing MAG biogeography.** When a query genome is searched against the GROS database, the closest MAG match in the database when at an ANI level >95% (same genomespecies) along with the physicochemical data of the sample from which the MAG was recovered and relative abundance in other samples will be shown on the interactive map and the left panel, respectively. The specific example shown represents the distribution of an unclassified *Oceanospirallales* MAG detected in the deep-sea plume during the DWH oil-spill.

The reference database consists of 2021 MAGs and SAGs, 1864 of which are of high quality (i.e., >75% completeness, <5% contamination) that were previously made available or were recovered from available metagenomes using established assembly and binning techniques^15^ as part of this study. For the latter, an iterative binning methodology^16^ was employed in certain cases where multiple metagenomes from a time or spatial series were available to recover additional high quality MAGs (mainly for the DWH impacted sediments); otherwise individual metagenomes were assembled and binned. The MAGs are grouped into species-like clusters based on the 95% genome-aggregate average nucleotide identity (gANI) threshold as recommended previously and implemented in MiGA ^17^, and only one representative genome for each cluster is used when a genome is queried against the database in order to reduce the CPU demand. A total of 1536 unique clusters (inter-cluster ANI < 95%) were recovered among these MAGs, including 22 clades with 5 or more members, thus revealing extensive species diversity for microbial taxa associated with oiled environments. Cultured taxa were observed in (only) 4.25% of all the recovered 95% ANI clusters (Fig. 2). The 2021 MAGs were assignable to 63 different bacterial and archaeal phyla based on genome-aggregate amino acid identity (gAAI) values (Fig. 3), with over 40 of them affiliated with Candidate Phyla Radiation (CPR) as well as the newly described Asgard superphylum ^18,19^. Recent studies have shown that Asgard archaea have the metabolic potential to carry out the anaerobic oxidation of short chain hydrocarbons ^18^. Moreover, notably, nearly 45% of the DWH MAGs had AAI values between 40-50% to the closest described species with available (isolate) genome(s), which represents at the least novel families ^14^ and further underscores the novel microbial diversity that exists in these oiled niches.

**Figure 2:**
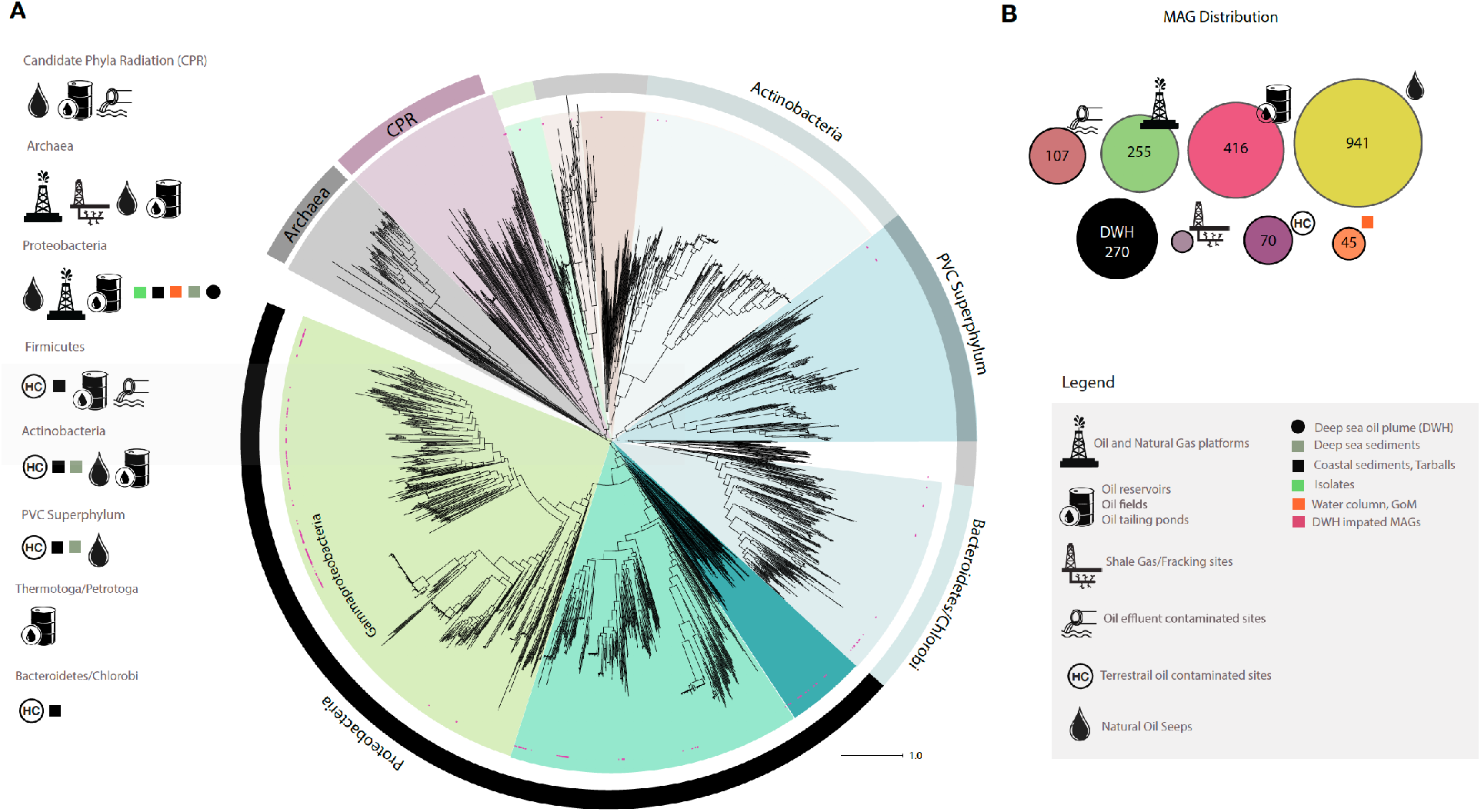
Whole-genome phylogeny of the oil-associated MAGs. **A**. The universal core gene phylogenetic tree of the1864 high-quality, oil-associated MAGs, when overlaid with the habitat of origin of the MAGs, reveals strong ecological niche specialization. For instance, proteobacterial MAGs appear widespread and not localized to a certain niche. However, MAGs from the CPR and Asgard superphyla were only associated with oil reservoirs, tailing ponds and deep-sea sediment in the vicinity of natural seeps. Similarly, MAGs from shale or natural gas fracking sites were either archaea or belonged to *Firmicutes* (specifically, the recently described “*Ca*. Frackibacter”). MAGs from *Thermotoga* or *Petrotoga* (extremophiles) were exclusively found in oil reservoirs or oil wells. Note that most of the MAGs recovered from DWH impacted sites (indicated as red dots in the tree) were assignable to *Proteobacteria* or *Bacteroidetes*. **B.** Total MAGs recovered from each ecological niche. Ecological niches are defined in the figure legend.

**Fig.3:**
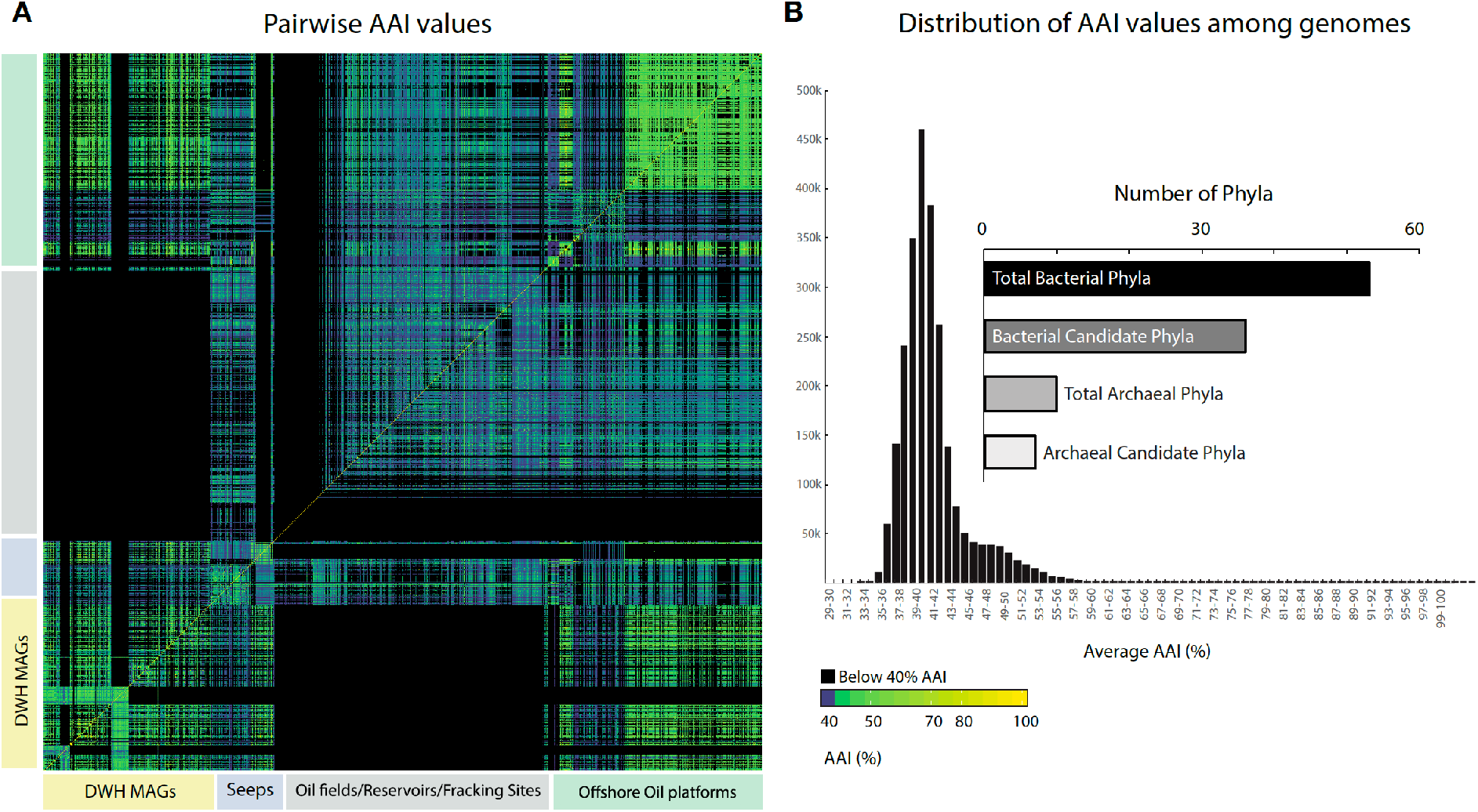
A. Pairwise genetic relatedness among MAGs based on AAI. Black boxes represent AAI values below 40%, blue boxes represent AAI values between 40-45%. **B.** Distribution of AAI values among the MAGs. Inset: Total number of bacterial and archaeal phyla represented by these MAGs.

The universal gene phylogeny revealed a strong ecological niche specialization for the majority of the MAGs and rather limited geographical distribution with a few exceptions. For instance, 82 % of the high quality DWH MAGs and SAGs were strictly associated with specific habitats of the Gulf, i.e., deep sea surface and subsurface sediments, deep sea oil plume or sediments and surface residue balls (or SRBs) recovered from coastal ecosystems, and they rarely crossed between these distinct habitats. Interestingly, at the 95% ANI (or species level), lab incubation studies that aimed to mimic the *in-situ* conditions after the DWH spill shared between ~18% and over 60% of the MAGs recovered from *in-situ* samples for the deep-sea plume ^20^ and coastal beach sands ^13^, respectively, which represents a much higher fraction compared to similar laboratory incubations for other microbial processes ^21,22^. At higher taxonomic levels (i.e., genus or higher), more overlap was observed between the lab simulations and respective field data (>80% overlap at the genus level) as well as between distinct habitats (~30% overlap) affected by the DWH spill.

MAGs associated with the DWH spill (across all habitats) predominantly belonged to the *Proteobacteria* phylum whereas a higher proportion of MAGs from sediments associated with natural oil seepage environments, petroleum reservoirs, and shale gas/fracking sites belonged to the recently proposed Candidate phyla radiation or CPR ^7^. Furthermore, archaeal MAGs were only found in the vicinity of natural deep-sea seeps, oil reservoirs and shale gas/fracking sites and were not detected in any of the DWH impacted samples. This is likely attributable to the lack of dissolved oxygen and the presence of gaseous hydrocarbons at the latter sites when compared to the DWH-impacted sites, most of which were aerobic. Archaea are typically associated with the presence of methane and other short-chain and gaseous hydrocarbons as is the case in natural oil seeps where dissolved oxygen is often not detectable ^18^. In contrast, the deep sea DWH plume at the seafloor was aerobic, presumably leading to enrichment of taxa that differed from those in the sub-surface samples. Further, the early stage composition of the deep sea DWH plume was rich in gaseous hydrocarbons, and as previous reports suggested, these gaseous compounds were instrumental in priming the microbial populations for breakdown of the more complex substrates of the plume ^23^. GROS can help in advancing such discoveries by documenting the abundances of MAGs along with the associated hydrocarbon and physicochemical gradient data and providing insights into the biodegradation process based on the MAGs’ successional patterns.

Mapping of metagenomic reads to the recovered MAGs was employed to determine the degree of biogeography (i.e., limited distribution globally) of the MAGs. For this analysis, reads from each metagenome were mapped against the MAGs at high nucleotide stringency (>95% nucleotide identity) and a non-zero TAD80 (*i.e.*, Truncated Average sequencing Depth across the genome after removing the top and bottom 10% genome positions in terms of sequencing depth) was used as the threshold to determine presence vs. absence of a MAG in the metagenome. Note that non-zero TAD80 corresponds to at least 10% sequencing breadth, corresponding to a confident detection (presence) at species-level resolution (ANI ≥ 95%) as suggested previously ^24^. This abundance information is also available through GROS for the reference MAGs of each 95% ANI cluster. In general, MAGs showed strong biogeography with the notable exception of the deep-sea oil plume and sea floor MAGs that showed broad distribution across geographic distances, even at the species level. More specifically, ~16% and ~17% of the MAGs recovered from the deep-sea oil plume and sediments belonging primarily to class Deltaproteobacteria and Dehalococcoidia were detected (i.e., same species) in the metagenomes recovered from the natural seeps (GoM, Santa Barbara Channel, Guaymas Basin) and seawater in the vicinity of oil platforms (Canada), respectively. There was considerable overlap between the beach sand and SRB communities across distinct geographic locations in the GoM (Figs.4A-C). Further, less than 10% of the coastal sediment associated MAGs (mostly proteobacterial) were observed in terrestrial crude-oil impacted metagenomes and typically at low abundances (Fig. 4D). Finally, the global marine metagenome data of the TARA Oceans expedition (exclusively uncontaminated samples) were only recruited by the uncontaminated GoM MAGs and, to a lesser extent, by the genomes of known hydrocarbonoclastic bacteria namely, *Alcanivorax spp*. and *Acinetobacter spp*. This finding could indicate that these hydrocarbon-degrading bacteria maintain substantial (not rare) populations in-situ based on growth on biogenic alkanes of algal or cyanobacterial origin, which are prevalent in the open, non-oil-contaminated ocean ^25^. In contrast, at least 40% of the MAGs recovered from the DWH oil plume and seafloor sediment impacted by the oil spill were not detected in the TARA ocean datasets (rare biosphere) but were found in low abundances in the metagenomes from the uncontaminated water column in the GoM. Collectively, these results could indicate that certain populations respond to oiling, i.e., increase from undetectable or low abundance to represent a substantial faction of the microbial community, typically >0.5-1% of the total upon oiling and are part of the rare biosphere in uncontaminated sites, but tend to be more abundant in the GoM due to abundant natural seeps in this region. In other words, the GoM appears to be more “primed” for crude-oil biodegradation based on our preliminary data.

**Fig. 4:**
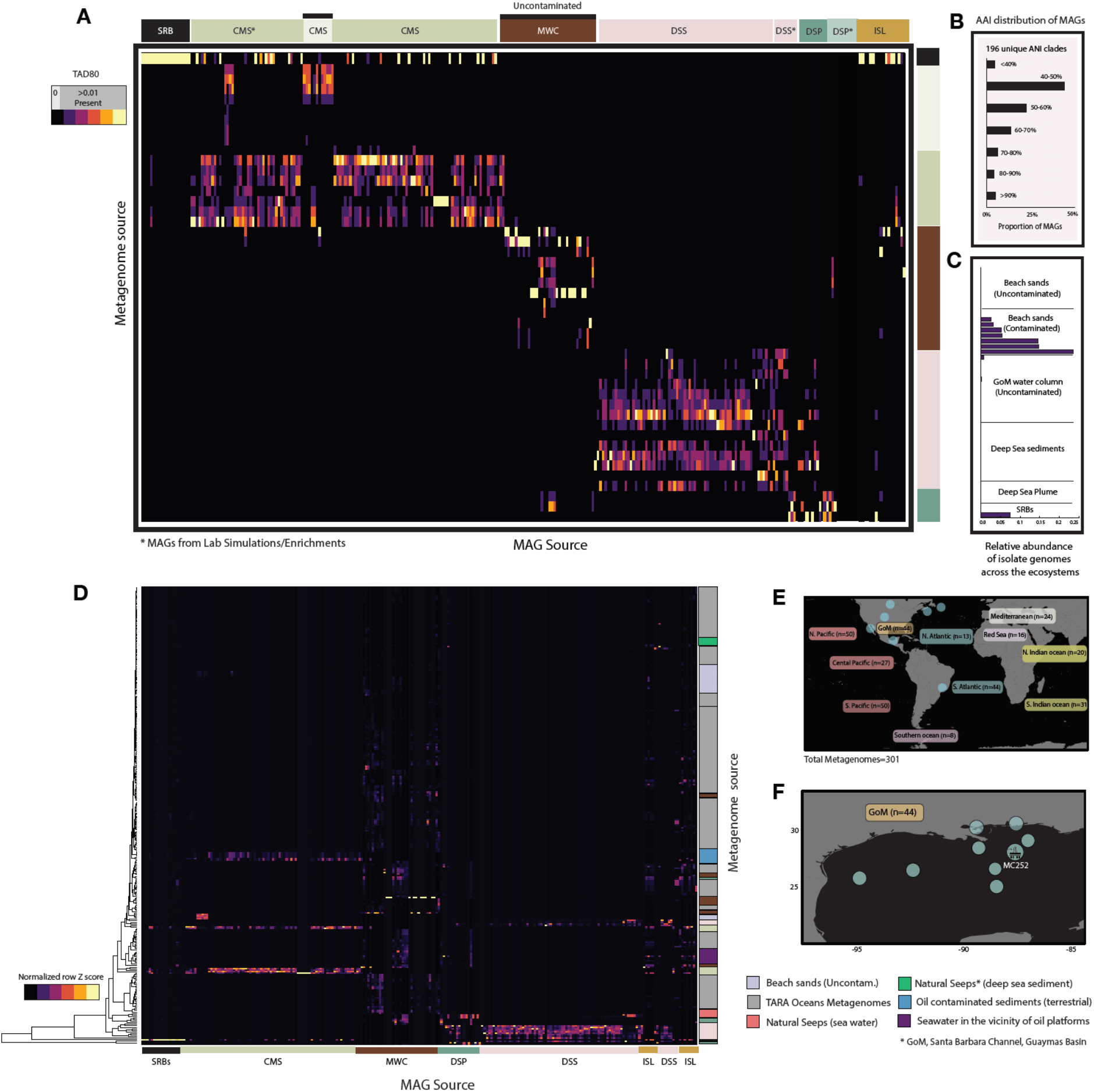
Biogeographic distribution of the DWH-associated MAGs. **A:** Heatmap showing the TAD80 values of DWH-associated MAGs in various metagenomes from the GoM (top x-axis); absence of a MAG in a metagenomic dataset (y-axis) is indicated in black color. S1-S4 represent lab simulation or enrichment data; uncontaminated water column samples were also included for comparison. Note the minimal overlap that occurs between habitats with the exception of the deep-sea sediment and plume. Substantial overlap also exists between the beach sand and SRB communities. **B.** Distribution of AAI values between the DWH associated MAGs and their closest described relative in the public databases showing that ~50% of the MAGs represent novel family or higher taxonomic ranks (e.g., best match AAI <50%). **C.** Proportion of reads from the metagenomic datasets mapping to isolate genomes of oil degraders showing that less than 5% of the reads mapped to these genomes. The notable exception is beach sands and SRBs where the recently described isolate “*Ca.* Macondimonas” made up >25% of the total community after the oil spill. The relative abundance for each dataset was normalized by their corresponding *rpoB* gene sequencing depth as described in the Methods section. **D.** Similar to Panel A but including 301 non-GoM metagenomes from uncontaminated global water column (includes TARA Oceans data), deep-sea natural seeps data from the GoM, the Santa Barbara channel and Guaymas Basin, crude-oil contaminated sediments and seawater data in the vicinity of oil and natural gas platforms, and oiled terrestrial (soil) metagenomes. Note that some of the uncontaminated water column samples shown (brown for GoM, grey for others) have very low or undetectable abundances for most of the oil MAGs, reflecting that they are members of the rare biosphere especially in non-GoM samples. **E**. Map showing the locations of the metagenomes used. **F.** Similar to panel E but showing only the GoM sampling locations. For simplicity, the habitats have been broadly categorized as Deep-sea sediment (DSS), Deep-sea oil plume (DSP), Coastal marine sediments (CMS), Surface residue balls (SRB) and marine water column (MWC) for uncontaminated samples recovered from the GoM.

In contrast to the strong biogeographical patterns largely observed at the individual MAG level, functional annotation of the metagenome reads and the MAGs associated with DWH impacted regions showed much less separation by habitat at hierarchical or subsystem levels (Suppl.Fig. 1A), i.e. specific functions did not show biogeographical or habitat-specific associations but the genomes that encoded them did. At a 95% ANI (or 95% nucleotide identity for individual genes), the DWH MAGs were grouped into 195 unique clusters whose MAG composition was highly dependent on the source habitat of the MAGs. However, when the proteins predicted from these MAGs were clustered at a 40% amino acid identity (subsystem level classification), no such clear clustering pattern was observed (Suppl.Fig. 1A, B). Moreover, the proteins involved in oil degradation (specifically, alkane monooxygenase as well as ring-hydroxylating dioxygenases involved in degradation of the aromatic hydrocarbons) did not show habitat preference indicating that these functions may have moved horizontally in the recent or more distant past between habitats (Suppl.Fig. 1A, 1C). Incomplete lineage sorting and secondary gene losses could also be other possible mechanisms underlying this observed pattern ^26^.

Despite recent efforts, many sites, pristine or contaminated, are under-sampled and additional environmental surveys are needed for a complete picture of oil-associated microbial diversity to emerge. The GROS project provides an easy integration of such data and capabilities to analyze new and unpublished data in the context of previously published data. This repository can be used to obtain a more holistic view of the microbial responses to oil, differences and similarities across habitats, and identify biomarkers that can be universally representative of the different phases of oil biodegradation and ecosystem recovery as exemplified above. The curated database may serve as a model approach for building similar resources for additional environmental processes and data of interest such as the waste/drinking water microbiome ^27^. GROS is publicly available through the link below and allows the user to download all MAGs and metadata mentioned above: http://microbial-genomes.org/projects/CrudeOilMicrobiome

## METHODS

### Data Curation

All publicly available sequence data at the time of this writing including shotgun metagenome and single-cell as well as isolate genome sequence data associated with the DWH oil spill that includes laboratory enrichments or simulations were used as a part of this study. Metagenome data from uncontaminated water column samples from the GoM were included to elucidate the baseline microbial data and assess the abundance of oil-associated MAGs/SAGs in such samples. To expand this dataset, data from all publicly available crude- or refined-oil contaminated ecosystems including, but not restricted to, natural oil seeps, petroleum reservoirs, coal fields, hydrocarbon contaminated sites, and methane gas vents were also included. All metagenomes and MAGs and SAGs were obtained from the NCBI or MG-RAST databases ^28^. Details on the data sources are provided in Supplemental Table 1.

### Quality Control and Trimming

The raw metagenome reads from the publicly available data (single or paired end) were trimmed using Trimmomatic (dynamic trim option) ^29^ and quality checked using the SolexaQA ^30^ package with a cutoff of Q>20 (>99% accuracy per base-position) and a minimum trimmed length of 50 bp. For certain metagenome datasets that had shorter average sequencing read lengths (~75 bp), a minimum read length of 30 bp was used as the threshold.

### Assembly and Binning

Assembly and binning were carried out for the metagenomes that did not have associated MAG data publicly available. Assembly (or co-assembly for metagenomes similar in composition) for the single-cell and metagenomic datasets was performed using IDBA-UD with default parameters unless otherwise noted ^31^ and only contigs >1000 bp were retained for binning. For time- or spatial-series metagenomic datasets (mainly sediment samples), an iterative subtractive binning pipeline was employed to enhance recovery of high-quality MAGs. In order to determine which of the metagenome sample sets could be co-assembled for the iterative binning, MASH, a tool employing the MinHash dimensionality reduction technique ^32^ was used to evaluate the pairwise distances between the metagenomic datasets. The resulting distance matrix was then visualized using NMDS (non-metric multidimensional scaling) plots. A combination of MASH distances and a Markov Cluster Algorithm (MCL) using the script “ogs.mcl.rb” from the Enveomics toolkit ^33^ that identifies Orthology Groups (OGs) in Reciprocal Best Matches (RBM), was employed to evaluate the samples that could be pooled together for co-assembly; samples with distance values below 0.1 were co-assembled. For the sediment metagenome samples alone, the paired end reads were merged using PEAR ^34^ with –p 0.0001 setting. The resulting reads (non-merged forward, non-merged reverse and merged) were combined and quality trimmed as described in the earlier section and sequences less than 70 bp were discarded. IDBA-UD with the following parameters: --long, --mink 25, maxk 121, --step 4 was used to assemble the resulting reads. These parameters provided longer contigs as well as improved recovery rates of MAGs from these environments.

Population genome binning was carried out using MaxBin v2.0 ^35^ and the MAG’s completeness and contamination were assessed via CheckM ^36^. MAG quality was determined as [Completeness – 5*(Contamination)]. All MAGs with quality score >50 were used in downstream analyses. High quality MAGs were defined as Completeness > 75% and Contamination < 5%, and medium quality MAGs were defined by Completeness >50% and Contamination > 10%. To determine redundancy of the MAGs (when multiple methods were used to recover MAGs, MAGs were de-replicated using a 95% ANI level over at least 20% of the genome using FastANI v1.1 ^37^.

### Spatio-temporal abundance distribution of DWH impacted MAGs

In order to assess the biogeography of the DWH impacted MAGs within and across habitats, the short reads from contaminated or non-contaminated metagenomes were mapped onto the MAGs. For this, the MAG sequences were initially tagged and combined into a single Bowtie 2 database ^38^. The metagenomic reads were then mapped onto this database using a competitive search using Bowtie 2. The 80% central truncated average of the sequencing depth or the TAD80 value was estimated for the mapped metagenome reads as described elsewhere ^16^. The estimated sequencing depth was normalized using the sequencing depth of a single-copy gene *rpoB* as described in Tsementzi et al. 2019 ^16^, which provide the normalzied relative *in-situ* abundance estimate across datasets. Normalized MAG abundances were visualized using a Heatmap generated using the heatmap3 package in R (v3.4.0) ^39^.

### Taxonomic and functional annotation of MAGs

Genes were predicted for the MAGs using MetaGeneMark ^40^ and annotated using the curated SwissProt database as described elsewhere ^41^. The resulting annotations were filtered by their amino acid identity and alignment score and bitscore using as a minimum match >= 40% AAI, >= 70% alignment length, bitscore >= 60. The SwissProt database identifiers were mapped to their corresponding metabolic function based on the hierarchical classification subsystems (Level 1 of the SEED subsystem category) ^42^ in order to provide broad functional category information. Genes relevant to oil biodegradation were manually verified. Heatmaps depicting the functional gene content across samples were generated using the heatmap3 package in R (v3.4.0). The MAGs were taxonomically characterized based on whole genome AAI and ANI comparisons against the NCBI prokaryotic database using MiGA ^14^.

### Whole genome phylogeny of the oil-associated MAGs and SAGs

Genes were predicted for all of the 1864 high quality MAGs recovered from crude-oil contaminated environments using Prodigal v2.6.1 ^43^. Only the MAGs with over 75% completeness were used in the whole-genome phylogeny reconstruction. A set of 106 universally present “marker genes” were identified in these MAGs using the script ‘HMM.essential.rb’ available as a part of the enveomics toolkit. The resulting reads were then aligned using Clustal Omega v1.2.1 and concatenated using the script ‘Aln.cat.rb’, which removes any invariable sites.

The phylogenetic tree was constructed using RaxML using the PROTGAMMAAUTO option (1000 bootstraps). The resulting tree was visualized using iTOL ^44^.

Maps used in the webserver were created using ArcGIS® software by Esri. ArcGIS® and ArcMap^TM^ are the intellectual property of Esri and are used herein under license.

## ACKNOWLEDGEMENTS

This research was made possible by grants from The Gulf of Mexico Research Initiative (RFP V Grant No 321611-00 as well as grants to the C-IMAGE II, C-IMAGE III, and Deep-C consortia). Data are publicly available at http://microbial-genomes.org/. Associated NCBI accession numbers have also been provided where applicable. The authors would also like to thank Ramachandra Sivakumar form the Center for Spatial Planning Analytics and Visualization (CSPAV) at Georgia Institute of Technology for his valuable input with the incorporation of GIS layers in the webserver.

The authors declare no conflict of interest.

