## Supplemental Fig.S1 for "Genome Repository of Oiled Systems (GROS): an interactive and searchable database that expands the catalogued diversity of crude oil-associated microbes"

**Supplemental Figures**

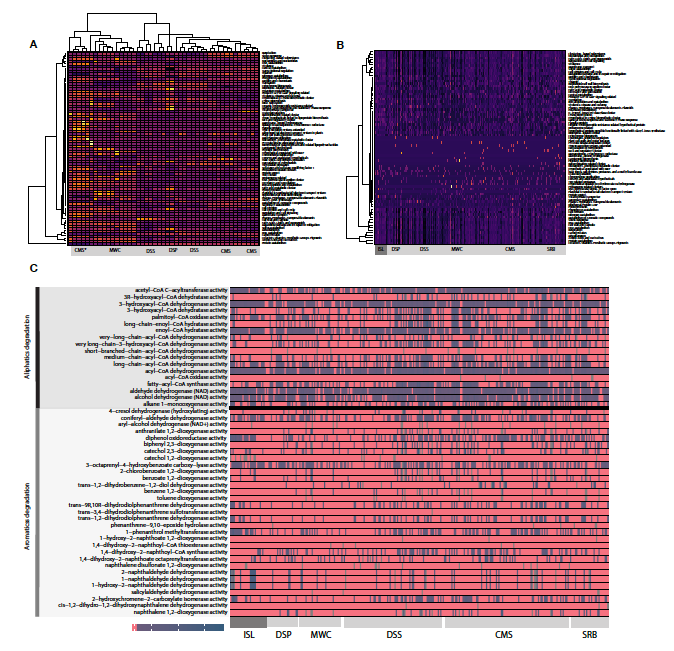

**Suppl. Figure 1: (Supplemental)** Heatmap showing SEED subsystem based clustering of the genes predicted in the metagenomes (A) and MAGs (B) clustered at 40% amino acid ID for the DWH impacted samples. C. Gene abundance for specific functions related to oil biodegradation detected in the MAGs. Note that no distinct clustering patterns are observed for the functional genes encoded by these MAGs. For simplicity the ecosystems have been broadly categorized as deep-sea sediment (DSS), deep-sea oil plume (DSP), coastal marine sediments (CMS), surface residue balls (SRB) and marine water column (MWC) for uncontaminated samples recovered from the Gulf of Mexico. Aerobic pathways have been indicated in black arrows and anaerobic pathways in red.
